# High-Resolution Phylogenetic and Population Genetic Analysis of Microbial Communities with RoC-ITS

**DOI:** 10.1101/2020.10.16.342691

**Authors:** Douglas B. Rusch, Jie Huang, Chris Hemmerich, Matthew W. Hahn

## Abstract

Microbial communities are inter-connected systems of incredible complexity and dynamism that play crucial roles in health, energy, and the environment. To better understand microbial communities and how they respond to change, it is important to know which microbes are present and their relative abundances at the greatest taxonomic resolution possible. Here, we describe a novel protocol (RoC-ITS) that uses the single-molecule Nanopore sequencing platform to assay the composition of microbial communities in unprecedented detail. This methodology produces long-read sequences including multiple copies of the same complete 16S ribosomal gene and its neighboring internally transcribed spacer (ITS) using rolling-circle amplification. The ribosomal 16S gene provides phylogenetic information down to the species-level, while the much less conserved ITS region contains strain-level information. When linked together, this combination of markers allows for the identification of individual ribosomal units within a specific organism, the assessment of their relative stoichiometry, and the ability to monitor subtle shifts in microbial community composition with a single generic assay. We applied RoC-ITS to a mock microbial community that was also sequenced using the Illumina platform, demonstrating its accuracy in quantifying the relative abundance and identity of each species.

## Introduction

The bacterial ribosomal RNA operon (rrn) is a polycistronic precursor gene containing the 5S, 16S, and 23S rRNA genes along with an internal transcribed spacer (ITS) region. After expression, the operon is cleaved to generate individual gene products, and the ITS region is degraded. The individual genes produce key RNA products that are involved in protein production and thus these genes, or their homologs, are found in all free-living organisms. Individual genomes may have one or more ribosomal operons (1) and the presence of multiple rrn copies is thought to allow cells to quickly respond by increasing growth rates in favorable environmental circumstances (2). Although multiple rrn copies are often homogenized by concerted evolution, there are many instances of intragenomic rRNA heterogeneity (1, 3).

The 16S gene has become the focus of modern microbial phylogenetics by virtue of its length and mix of highly conserved and variable regions. The conserved regions make it amenable for polymerase chain reaction (PCR) amplification with conserved primers, while the variable regions make it a useful phylogenetic marker (4). As a result, it has a long history of use in phylogenetics, and is widely used to taxonomically survey microbial populations (5, 6). While longer, the 23S gene has a lower density of informative markers, and is therefore rarely used for general phylogenetic purposes in eubacteria (4); the 5S gene is small and relatively rapidly evolving, and has seen occasional use as a phylogenetic marker (7–9). The ITS region is the most rapidly evolving part of the rrn, likely because it has no defined functional role (though occasionally one or more tRNA genes are found within it). This poor conservation and variability in length has made it difficult to sequence. However, the position of the ITS region between the highly conserved portions of the 16S and 23S genes means that it can be readily amplified with conserved primers. It has therefore been used for high-resolution DNA fingerprinting in a technique called ARISA (10–12).

Due to its mix of conserved and variable regions, the 16S gene has long been the primary target for phylogenetic and taxonomic study. The full-length 16S can be used for phylogenetic resolution down to the species level (13), and was readily acquired using Sanger sequencing. As 454 (14, 15) and later Illumina (16) short-read sequencing technologies became available at much lower costs per base, 16S sequencing shifted to focus on one to three of the nine recognized variable regions (14, 17). Depending on the organisms involved, this smaller number of regions was sufficient to classify microbes taxonomically to the family or genus level, but lacked the phylogenetic resolution of the full-length gene. On the other hand, metagenomic studies of random genomic regions have revealed tremendous genetic and functional diversity within species (18–20), indicating that short-read sequencing of 16S fails to reflect much of the within species variation. However, the low cost of short-read sequencing platforms resulted in an explosion of microbial data, along with new databases and tools for analyzing, profiling, and comparing these potentially enormous datasets (21, 22).

Recently, new developments in sequencing technologies have ushered in attempts to combine quantity with quality, and to return 16S sequencing to its former heyday. Long-read, single molecule sequencing methods including the Pacific Biosciences and Oxford Nanopore Technologies sequencing platforms have made it possible to sequence thousands of bases at a time (23–27). These sequencing techniques have a high error rate that keeps them from being useful for rrna profiling by themselves; however, creative use of long reads to repetitively sequence a circularized template result in “consensus reads” with reduced error rates (11, 28–32).

Here, we describe a new high-throughput sequencing strategy that relies on rolling-circle amplification coupled with Oxford Nanopore Technologies (hereafter “Nanopore”) long-read single molecule sequencing to capture the entire 16S and ITS region. By sequencing the entire 16S region our method provides high-quality phylogenetic information, including resolution below the species level. The inclusion of the ITS region also allows for the ability to distinguish among individual rrn copies within a genome. Together, these tools allow for resolution between microbial strains, leading to more complete characterization of microbial populations and their dynamics. We describe the steps necessary to prepare and sequence a library of reads using our method, which we call RoC-ITS (“rockets”), and detail a computational pipeline that can quickly and effectively analyze the data. We apply RoC-ITS to a mock community of eubacteria that have also been sequenced with Illumina short-reads in order to demonstrate its effectiveness.

## Results

### Sequencing of Mock Microbial Community

The 16S-ITS region was PCR amplified from a diverse stratified mock community of 20 different Bacteria (Table 4). The 16S-ITS product was circularized using Gibson assembly, converted into a single-stranded template using rolling circle amplification, and sequenced on a previously used Nanopore Minion flowcell (see Methods). In total, we generated 31,215 sequences, of which 25,489 passed the initial Nanopore quality filters (the reuse of a flowcell is expected to produce a much lower number of reads compared to a new flowcell). The phi29 polymerase is highly processive and produced single-stranded sequencing templates that were often 50 kb or longer from the circularized 16S-ITS product. Given that the 16S-ITS product should be between 2000-3500 bps in length, the rolling circle amplification should, in theory, have produced between 12-25 copies of the 16S-ITS region in each sequencing template. The mean Nanopore read length was 12.7 kb with a mode between 4-5 kb (Supplemental Fig 1). In parallel with the generation of the Nanopore data, we also generated a traditional V4 region dataset using Illumina sequencing. This dataset had a total of 286,793 paired-end reads that were generated on a MiSeq run.

Nanopore reads have a significant error rate, mostly in the form of insertions and deletions. These errors mean that RoC-ITS reads require further computational processing to produce a high-quality consensus sequence that is suitable for further analysis. The first step was to identify and isolate each of the sub-reads within a longer Nanopore sequencing read. To do this, we developed a DNA hidden Markov model (HMM) that tolerates a high rate of insertions and deletions. The HMM was designed to recognize the splint DNA and the flanking regions containing the unique molecular barcodes. After aligning the HMM to the Nanopore reads we extracted the 16S-ITS portion into sets of sub-reads. The bulk of the sub-reads (over 80%) were between 1800 and 3100 bp (Supplemental Fig 2), with most of the remaining sequences much smaller than would be expected. To ensure that we captured potentially interesting edge cases, we ultimately selected the 5,442 Nanopore reads with two or more sub-reads that were between 1500 and 3500 bp in length.

Another form of error was discovered at this point: some sets of sub-reads from the same RoC-ITS read had much higher sequence divergence from each other than expected. These distinct sub-reads showed a periodic pattern within the longer RoC-ITS read, consistent with the fusion of two or more 16S-ITS products into a single circular molecule (Supplemental Fig 3). These cases were processed to separate the intermingled sub-reads into distinct sets before further processing (see Supplemental Methods for details). There were 4,955 sets of sub-reads with a single distinct insert and 487 with two or more distinct inserts. In the end, each set of distinct sub-reads was processed to produce a consensus RoC-ITS sequence. To ensure that we had the highest quality reads for further analysis, we further filtered the RoC-ITS sequences retaining only the 2,569 generated from seven or more sub-reads. At this point the RoC-ITS sequences can be treated like any other PCR amplicon, including identifying and removing chimeric sequences. After this final filtering step, we had 1,668 high-quality consensus sequences (6.5% of original Nanopore reads).

### Phylogenetic Analysis and Classification of 16S-ITS Reads

The Illumina 16S V4 amplicon reads were quality-filtered and analyzed with the QIIME2 pipeline, resulting in 239,988 valid sequences. QIIME2 reported 18 different Amplicon Sequence Variants (ASVs) belonging to 11 different genera (Table 4). These ASV and taxonomic classifications were then used in the analysis of the 16S-ITS data (i.e. the Nanopore data) to provide a common framework in which to categorize the longer reads. The 16S-ITS sequences were also run through QIIME2 using only the same V4 region used in the Illumina data. This approach allowed us to classify 90% of the RoC-ITS sequences (N=1,504).

Using BLAST against the ASVs we were able to taxonomically bin all of the 16S-ITS reads and, when there were both BLAST and QIIME2 results available, no disagreements between these two approaches were found. The relative abundance of the different ASVs in the Illumina and Nanopore sequences were not statistically different (Table 4; p-value = 0.1567; (33). Some of the least abundant taxa (*Mycobacteria*, *Rhodococcus*, and *Inquilinus*) could be identified in the Nanopore reads initially but were lost during the various filtering and cleaning steps. The *Micrococcaceae* family seems to be the most different in frequency between datasets, with both the Bacilli and Pseudomonads showing small differences (Table 4). These effects do not have a clear correlation with 16S-ITS length and were present from the start of the analysis, so cannot be entirely attributed to a processing artifact. More likely these differences reflect differences in the efficacy of the primers between the V4 and 16S-ITS regions for these particular organisms; such issues can be explored and potentially improved upon with better primers in the future.

We generated phylogenetic trees from the 16S-ITS sequences to assess the variation present in the data. The ITS portion of the 16S-ITS consensus reads were too diverged between taxa to generate an MSA for phylogenetic analysis across all species. Therefore, we initially extracted and generated the MSA from only the 16S portion of the 16S-ITS sequences. A phylogenetic tree was generated from the MSA using RAxML (Fig 2). This tree is highly congruent with the accepted relationships among families included in the mock community.

The 16S tree also shows the level of divergence between sequences within families (Fig 2). Using only the 16S gene, an individual read was typically 2-3% different from the consensus sequence for that ASV group (Table 4). This error rate is higher than would be expected for Illumina-based sequencing. The cause of this variation reflects a combination of factors, including the actual differences between the 16S genes within a given organism (see next section), the errors associated with Nanopore sequencing (especially with a re-used Nanopore flowcell), errors introduced during processing, and any errors introduced and accumulated during the PCR steps. Additionally, we cannot discount the possibility that there were chimeras that were either too small or so similar in sequence—e.g. between two different 16S genes from the same species—that they were not detected, and thus may have driven up the error rate. While this error rate is high, it does not mask the underlying taxonomic signal and still allows for the identification of individual 16S genes within a species.

### Detecting multiple ribosomal operons

Many organisms contain multiple rrns that are often difficult to distinguish based on the sequence of the 16S gene alone, however the different operons may be identifiable based on the differences within the ITS because of its generally low conservation. Being able to distinguish and count individual rrns in a genome would produce better abundance and diversity estimates for a microbial community (34, 35). As an example, we built a phylogenetic tree of the 16S-ITS sequences for the *Duganella* isolate in the mock community and identified six distinct 16S-ITS regions with at least moderate bootstrap support (70%+) (Fig 3). These different clades exemplify the power of RoC-ITS: a tree of the *Duganella* 16S gene alone is not able to identify distinct clades with high bootstrap support (Supplemental Fig 4). This result indicates that the 16S-ITS sequences are more powerful tools for distinguishing intra-genomic rrn variation.

When constructing the *Duganella* 16S-ITS tree we included the 16S-ITS regions from two complete *Duganella* genomes (Fig 3). The complete genomes each possess 7 rrns, while the clustering of the *Duganella* RoC-ITS reads identified 6 distinct clades. Interestingly, the largest clade (Group 0 in Fig 3a) contains 23 16S-ITS reads, which is nearly double the average of the other 5 clades (10.8, s.d. = 3.1). which itself is close to the mean expected if there were 7 rrns in which case we would expect 11.6 sequences per clade. That makes Group 0 nearly four standard deviations from the mean of the other clades, a value that is highly statistically significant (p-value = 3.5e^−5^). Even including Group 0 in the mean and standard deviation calculation, Group 0 is still significantly different from the mean (p-value = 0.026). All of this suggests that Group 0 likely represents two distinct rrns, possibly missed in the phylogenetic analysis because of a recent gene conversion event. If there are indeed 7 rrns, then this result provides further support for the idea that the RoC-ITS approach is quantitative and can be used to estimate the number of rrns in a genome.

## Discussion

In this work, we describe an approach using Nanopore sequencing to capture the entire 16S gene and ITS regions, demonstrating that these long, variable-length sequences can be effectively and quantitatively sequenced. While 16S sequencing has been frequently used to assess microbial community structure, the particular regions examined are largely dictated by the sequencing technologies applied. Sanger sequencing is low-throughput and relatively expensive but can generate high-quality, full-length 16S gene products that are amenable for phylogenetic analysis down to the species level. Illumina sequencing strategies produce much larger numbers of short reads, and are used to assay the more variable sub-segments of the 16S gene; this approach allows for taxonomic profiling of even complex microbial communities down to the genus level.

Our method, which uses rolling circle amplification to sequence each 16S-ITS region multiple times, has the ability to identify different ribosomal operons within a single genome and to measure their relative abundance. By including ITS with the already widely used variable regions of 16S, our method also has the ability us to distinguish microbial relationships at finer resolutions. The addition of the ITS sequence and copy-number information should allow for more nuanced classification, and can distinguish microbes down to the strain-level. As we showed with paired sequencing of a mock microbial community using RoC-ITS and Illumina V4, these two approaches were highly congruent with no evidence that the variable length of the ITS region impacted the outcome.

Our approach could easily be made applicable to a broad range of organisms. With minor modifications to the primers or a multiplex PCR strategy, both archaeal as well as eubacterial genome 16S-ITS regions could be amplified. For fungi, ITS-23S primers should allow similarly long sequences and phylogenetically rich datasets. Together, this should give us a much richer and detailed view into microbial communities across a broad spectrum of microbes. For these things to become possible, the sequencing strategy proposed here (or a similar strategy) would need to be much more widely adopted. Such an overhaul would also replace and expand upon databases full of 16S variable region data (e.g. SILVA; 35).

While promising, the technique introduced here can still produce erroneous sequences. As with any PCR-based strategy, the long, highly similar templates generated by RoC-ITS are prone to chimerism, which can confound the identification and classification of specific reads. The use of Gibson assembly can also introduce a kind of chimerism at the level of Nanopore reads, with hetero-concatemeric circular templates that complicate consensus generation. Large databases of trusted 16S-ITS sequences would allow existing chimera detection software to run more effectively, though these should be modified to work with new error models based on Nanopore consensus sequences. Bottlenecking of the sample, along with the use of molecular barcodes, could also help to identify and potentially eliminate chimeras. Identification of the hetero-concatemeric templates and how best to resolve these will require additional and larger studies looking at whether these sequences behave differently from the individually circularized templates. Choosing the right sequencing strategy is key to generating high-quality consensus sequences. Our use of rolling circle amplification to repeatedly sequence the same 16S-ITS gene resulted in significant improvement in sequence quality, and this approach is likely to improve as the Nanopore sequencing technology matures. However, our approach did require considerable computational effort to generate high-quality sequences and led to a ^~^15-fold reduction in output. Future improvements to the computational pipeline may also improve yield.

Though we used the Nanopore platform to generate the 16S-ITS reads here, the same approach should be achievable using a variety of different sequencing technologies. For instance, 16S-ITS sequences could also be iteratively sequenced using the PacBio platform. However, this approach will also require significant computational effort to generate the consensus, and might result in biased amplification due to size variation in the ITS region (although the sequence-level errors may be reduced). Using Illumina sequence, the loop-seq sequencing strategy (36) may also be compatible with sequencing of these 16S-ITS products, at least given the amplicon lengths seen here. In loop-seq, the amplicon sequence is cloned into a molecularly barcoded vector, amplified, and then random deletions are introduced specifically in the amplicon sequence. This allows for coverage along the length of the amplicon to be recovered along with the molecular barcode. This information can then be used to assemble and reconstruct the original amplicon sequence. While such an approach would require at least a 300-cycle Illumina sequencing run, it may result in higher quality products given the underlying high-quality reads. Eventually, given a well-characterized database of ITS sequences, it may eventually be possible to dispense with the 16S portion altogether and rely specifically on the ITS region for classification.

In addition to different sequencing technologies, there are other modifications that could improve or expand on the approach outlined here. For instance, RoC-ITS could be applied to RNA data in order to quantify gene expression among different rrns. While it is unclear whether this approach would capture expressed rrns due to the degradation of the ITS, it may be that a 16S-ITS approach would provide an accurate instantaneous measure of rrn expression, albeit one that is organism dependent. Finally, while we have emphasized the use of this approach for the 16S and ITS loci, this approach is perfectly amenable to any other large locus that needs to be sequenced in a targeted manner.

## Methods

The RoC-ITS approach (Fig 1) involves multiple amplification, circularization, and linearization steps. We describe each of these in turn, followed by additional methods used in this study. Table 1 contains a glossary defining several terms commonly used in the text.

**Figure 1.**
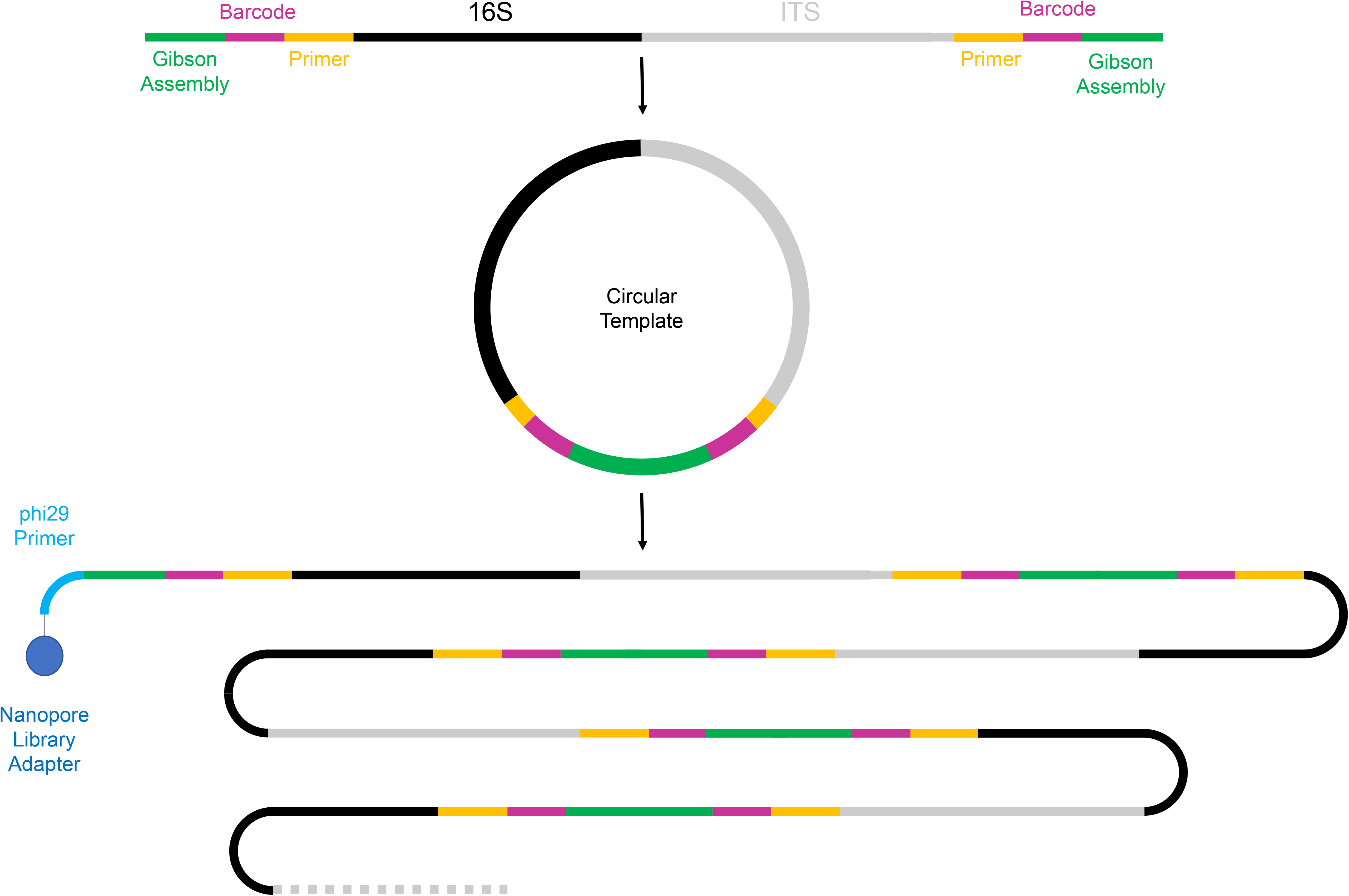
Diagram illustrating the RoC-ITS strategy. The 16S-ITS region is isolated using PCR and Gibson Assembly is used to circularize the products. A long single-stranded DNA product consisting of multiple iterations of the same 16S-ITS region is generated using the phi29 polymerase. Finally, the linear product is used to create an Oxford Nanopore DNA library for sequencing.

**Figure 2.**
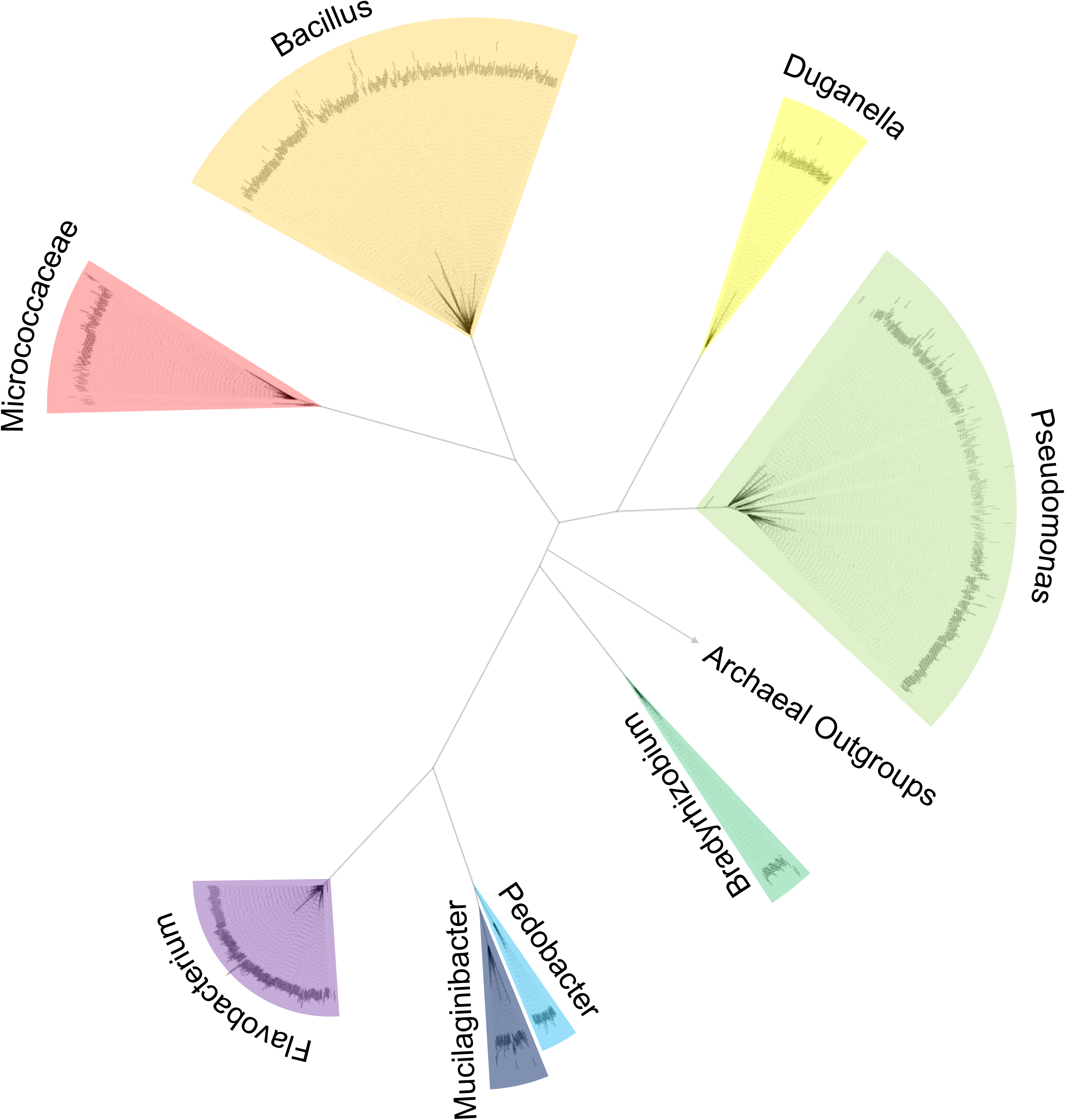
Phylogenetic tree of the RoC-ITS 16S regions from the mock community, using archaeal sequences as an outgroup. The multiple sequence alignment was structurally informed to allow comparison of the eubacterial and archaeal sequences and the tree was generated with RAxML using 100 bootstraps. The major eubacterial lineages from the mock community have been highlighted with wedges. All labeled branches have 100% bootstrap support; the branch to the archaeal outgroup has been shortened for clarity.

**Figure 3.**
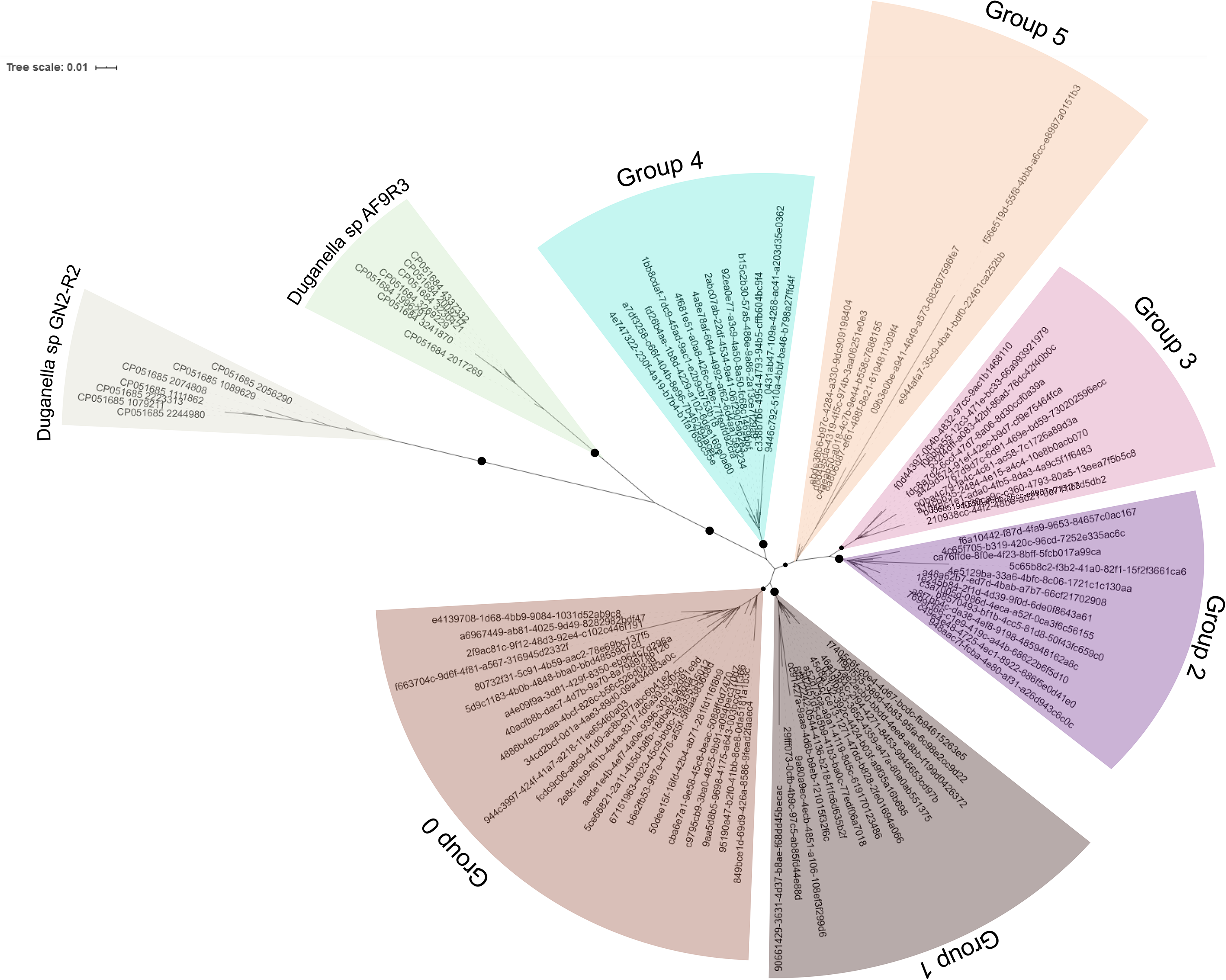
Phylogenetic tree generated from *Duganella* 16S-ITS sequences using RAxML with 100 bootstraps. Seven 16S-ITS regions each were identified and extracted from two complete *Duganella* genomes, *Duganella* sp GN2-R2 and *Duganella* sp AF9R3 (accessions CP051685.1 and CP051684.1 respectively) and included in the analysis; their individual 16S-ITS sequences act as outgroups to the Nanopore derived RoC-ITS sequences derived from the *Duganella* included in the mock community. The individual clades of have been highlighted with wedges and labeled. Bootstrap values for important branches have been indicated with filled circles on the applicable branch. Large circles indicate 95% or greater bootstrap support while smaller circles indicate 70% or greater support.

**Table 1.** Glossary of Commonly Used Terms.

| Term | Definition |
| --- | --- |
| 16S-ITS Region | An instance of the 16S ribosomal RNA and its adjacent ITS region |
| 16S-ITS Product | The PCR product consisting of an instance of the 16S and ITS region defined by the RoC-ITS 27F and RoC-ITS 189R primers |
| V4 Region | The 4th variable region of the 16S gene as defined by the Earth Microbiome F515/R806 primers |
| RoC-ITS Consensus Sequence | Consensus sequence derived from multiple independent RoC-ITS sequences |
| RoC-ITS Sequence | Consensus sequence derived from the amplification and iterated sequencing of a specific 16S-ITS region |
| RoC-ITS Read | A long Nanopore read derived from a circularized 16S-ITS product consisting of 0 or more sub-reads |
| Sub-Read | An individual 16S-ITS product found in a RoC-ITS read |
| Splint DNA | Derived from the lambda virus genome, a sequence used to circularize the 16S-ITS products |
| ITS | Internally transcribed spacer region; part of the ribosomal RNA operon |
| ASV | Amplicon sequencing variants; unique sequences identified by DADA2 here derived from the V4 region |
| Set of Sub-Reads | Multiple sub-reads derived from a single RoC-ITS read |
| MSA | Multiple sequence alignment |
| rrn | Ribosomal RNA operon including the 16S and 23S genes flanking the ITS region |

### RoC-ITS: PCR amplification

RoC-ITS uses PCR to amplify the 16S-ITS region from a DNA sample. The RoC-ITS 27F 16S primer (5’- <u>AATGATACGGCGACCACCGAGAT</u>NNNNNAGAGTTTGATCMTGGCTCAG-3’) was derived from the 27F 16S primer (Chen et al., 2015) while the RoC-ITS 189R 23S primer (5’- <u>ATGGAAGACGCCAAAAACATAAAGGCTGC</u>NNNNNTACTDAGATGTTTCASTTC-3’) was derived from the 189R primer (Hunt et al., 2006). In combination, this primer pair should amplify the 16S-ITS regions from eubacterial organisms. Both primers contain a 5-base molecular barcode and have a unique 5’ sequence (underlined; referred to as RoC-ITS 27F Unique and RoC-ITS 189R Unique, respectively). PCR reactions contain 20 ng input DNA in 12.5μl of KAPA 2X Master Mix (KAPA HiFi HotStart ReadyMix. Catalog number: KK2602. Vendor: Roche Diagnostic), 10μM forward primer and/or 10μM reverse primer, and nuclease-free water to bring the final volume to 25μl.

The overall PCR recipe is given in Table 2. Briefly, the first round of PCR adds a molecular barcode to the 16S side of the template. The 16S-ITS product is then cleaned with a 0.5x bead cleanup (HighPrepTM PCR paramagnetic bead solution. Vendor: MAGBIO. Catalog number: AC-60050 using a MagStrip Magnet Stand 10. Vendor: MAGBIO. Catalog number: MBMS-10) to remove any unused primer. The second round of PCR and cleanup similarly adds the molecular barcode on the 23S end of the molecule. Subsequent rounds of PCR then amplify only those products that possess both molecular barcodes. We examined the PCR product on an Agilent TapeStation using a D5000 tape to ensure the bulk of the product was between 2000 and 3500 bp in length.

**Table 2.** RoC-ITS PCR Program.

| Segment | Temp (C) | Time (sec) | Cycles | Primers |
| --- | --- | --- | --- | --- |
| Initial denaturation | 95 | 03:00 | 1 | RoC-ITS 27F 16S<br>only |
| Annealing | 46 | 00:20 |  |  |
| Extension | 72 | 05:00 |  |  |
| Cleanup |  |  |  |  |
| Denaturation | 95 | 03:00 | 1 | RoC-ITS 189r 23S<br>only |
| Annealing | 46 | 00:20 |  |  |
| Extension | 72 | 05:00 |  |  |
| Cleanup |  |  |  |  |
| Initial denaturation | 95 | 03:00 | 1 | RoC-ITS 27F Unique<br>+<br>RoC-ITS 189r Unique |
| Denaturation | 98 | 00:20 | 12 |  |
| Annealing | 60 | 00:20 |  |  |
| Extension | 72 | 05:00 |  |  |
| Final extension | 72 | 10:00 | 1 |  |
| Hold | 4 | 00:00 | 1 |  |
| Cleanup |  |  |  |  |

### RoC-ITS: Splint Amplification

A splint molecule containing part of the lambda phage genome was generated via PCR similarly to the protocol of Volden et al. (37). This splint was 386 bp long and has sequence identity with the RoC-ITS 27F Unique and RoC-ITS 189R Unique primers. The splint was generated by PCR with two primers, Lambda_F_23S <u>CTTTATGTTTTTGGCGTCTTCCAT</u>AAAGGGATATTTTCGATCGCTTG, where the underlined portion matches the RoC-ITS 189R Unique primer, and the Lambda_R_16S <u>ATCTCGGTGGTCGCCGTATCATTT</u>GAGGCTGATGAGTTCCATATTTG, that matches the RoC-ITS 27F Unique primer. The PCR was carried out with 1 μl of Lambda DNA (20 ng/μl; Catalog number: N3011S. Vendor: New England Biolabs Inc), 12.5 μl of KAPA 2X Master Mix, 1 μl of 10 μM Lambda_F_23S and the Lambda_R_16S primers, and nuclease-free water to bring it to its final volume (25 μl). The PCR recipe for this step is shown in Table 3. Following PCR, the product was eluted in 30 μL of nuclease-free water after a 1.2x bead cleanup.

**Table 3.**
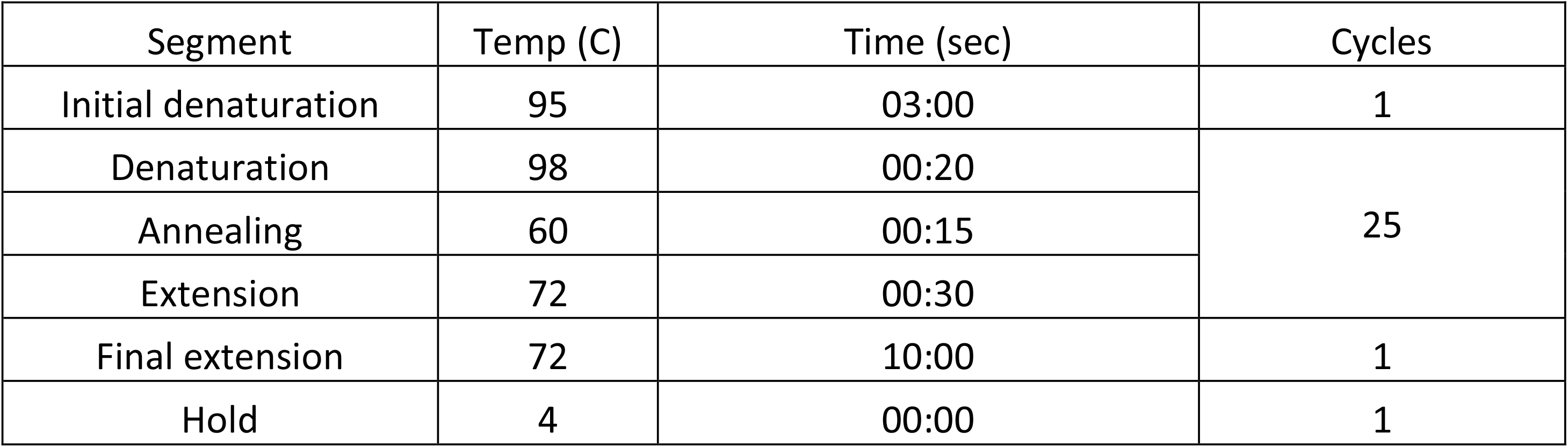
Splint PCR Program.

| Segment | Temp (C) | Time (sec) | Cycles |
| --- | --- | --- | --- |
| Initial denaturation | 95 | 03:00 | 1 |
| Denaturation | 98 | 00:20 | 25 |
| Annealing | 60 | 00:15 |  |
| Extension | 72 | 00:30 |  |
| Final extension | 72 | 10:00 | 1 |
| Hold | 4 | 00:00 | 1 |

### RoC-ITS: Circularization

The linear RoC-ITS product is circularized with a DNA splint that matches the RoC-ITS 27F Unique and RoC-ITS 189r Unique primers using Gibson assembly (Gibson et al., 2009). In a 20 μl reaction, 150 ng of RoC-ITS product and splint DNA are combined with 10μl of Gibson Assembly 2X Master mix (Catalog number: E2611S Vendor: New England Biolabs Inc), and are incubated at 50°C for 60 minutes. The reaction is bead-cleaned with 1.0x bead ratio (HighPrepTM PCR paramagnetic bead solution. Vendor: MAGBIO. Catalog number: AC-60050) and eluted in 42 μl of nuclease-free water. Linear molecules in the reaction mixture are then degraded using an exonuclease with the reaction carried out in a 50 μl volume: 2 μl of 25 mM ATP solution, 5 μl of 10X reaction buffer and 1 μl of Plasmid-Safe DNASe at 37°C for 30 minutes. The circularized reaction product is isolated with a 1.0x bead cleanup and eluted in 12 μl of nuclease-free water.

### RoC-ITS: Rolling Circle Linearization

The circularized RoC-ITS product is converted into a long single stranded linear product using the phi29 polymerase (Fig 1). The phi29 initiation primer (5’- <u>CGCCAGGGTTTTCCCAGTCACGAC</u>GAAGACGCCAAAAACATAAAG-3’) has a unique 5’ end (underlined) that does not match the circular template; it is used during the subsequent Nanopore library construction step. The 3’ end of the splint-primer was designed to anneal to the region adjoining the splint DNA and into the RoC-ITS 189r Unique primer, which should occur only once per circularized molecule (Fig 1). This means that only a single product is produced per circularized molecule, without the usual complications (i.e. branched products) associated with phi29 hexanucleotide-based amplification procedures (37). The length of the phi29 product is dependent on the stability and processivity of the polymerase and independent of the size of the plasmid, thereby removing the variable length of the ITS segment as a bias during later steps. As only a single priming site was present per molecule, no true amplification occurred except by the polymerase iteratively processing around the circle, such that the long product contains multiple instances of the PCR product. Production of the final linear product is performed in five parallel reactions to generate enough material for subsequent library production. Each reaction is performed in 50 μl at 30°C overnight with 2 μl of circulated DNA with 2.5 μL of 10 μM (each) dNTPs (Catalog number: N0446S Vendor: New England Biolabs Inc), 2.5 μL Rolling cycle primer (10 μM), 37 μL ultrapure water, 5 μL of 10× Phi29 Buffer, and 1 μL of Phi29 (Catalog number: M0269L Vendor: New England Biolabs Inc). The combined products are cleaned at a 0.5x bead ratio and eluted in 10 μl of nuclease-free water.

### RoC-ITS: Nanopore Library Construction and Sequencing

Oligos A (5’- GGCTTCTTCTTGCTCTTAGG-3’) and B (5’- GTCGTGACTGGGAAAACCCTGGCGCCTAAGAGCAAGAAGAAGCCA-3’) are designed to anneal to the phi29 initiation primer and to each other to generate an A-overhang required for the Nanopore library kit to function (Fig 1). This pair of primers is also used to dictate which strand of the library will be preferentially sequenced by the Nanopore flowcell. The annealing buffer (10 mM Tris-HCl pH7.5, 50mM NaCl) containing 1.4 μM each of oligoA and oligoB is heated to 95°C for 2 minutes and allowed to slowly cool to room temperature. The annealed oligoA/B product is then covalently linked to the rolling circle linear product with a ligation reaction: 9.5 μl of cleaned rolling circle product with 1 μL of annealed oligoA/B adaptor, 3.0 μL of NEBNext Quick Ligation Reaction Buffer and 1.5 μL of T4 DNA Ligase (NEBNext Quick Ligation Module; Catalog number: E6056. Vendor: New England Biolabs Inc) incubated at room temperature for 10 minutes. The oligoA/B adapters are removed from the product with a 1x bead cleanup and eluted into 60 μL of nuclease-free water. A Nanopore DNA library is generated using the standard kit protocol: 60 μL of cleaned adaptor ligated DNA with 25 μL of ligation buffer (LNB), 10.0 μL of NEBNext Quick T4 DNA ligase and 5μL of Adapter Mix (AMX) (LNB and AMX are provided in Nanopore SQK-LSK109) and incubated at room temperature for 10 minutes. The DNA library is purified with a 0.4x bead cleanup and eluted in 15 μL of elution buffer (from Nanopore SQK-LSK109 kit) and is designed to then run Nanopore SQK-LSK109 flowcell R9.4.1 following the prescribed Nanopore protocol. In the data shown here, we use a previously used but cleaned flowcell of this type. Base calling was performed using the Oxford Nanopore guppy_basecaller (version 3.4.4+a296acb) using a GP100GL video card (Tesla P100 PCIe 16GB, rev a1; NVIDIA Corporation, CA, USA) to generate the RoC-ITS reads.

### Processing RoC-ITS Reads

Individual Nanopore reads have a high error rate, with frequent insertions and deletions. To identify individual 16S-ITS products, we constructed a DNA HMM using the HMMer software (hmmsearch v3.2.1 with default parameters) to identify the known splint DNA and neighboring PCR primer sites along with the molecular barcodes (Eddy, 2011). A set of sub-reads for a given RoC-ITS read was defined as all the regions not identified by the splint DNA HMM that were between 1500 and 3500 bp in length. Each set of sub-reads from a single circle was then converted into a multiple sequence alignment using the prank alignment software (http://wasabiapp.org/software/prank/) resulting in a tree and multiple sequence alignment (MSA) file. The MSA was then further refined using probcons (version 1.1; (38) to resolve small issues with consistency in the prank MSA (see the supplemental document for additional details and an example). This improved MSA was then used to generate a consensus sequence, hereafter referred to as a RoC-ITS sequence, that can be analyzed further.

### RoC-ITS Reads with Multiple Distinct Sub-Reads

A small fraction of the RoC-ITS reads generated poor consensus sequences that did not match known or expected 16S genes. These appeared to be hetero-concatemers consisting of two or more distinct 16S-ITS PCR products combined into a single circular template during the Gibson Assembly step and read out in a single Nanopore read. The details of how to identify and resolve these situations are presented in the supplemental text.

### Consensus Generation from Multiple Sequence Alignments

A final RoC-ITS sequence was derived from the prank/probcons MSA of its sub-reads using a custom script (https://github.com/mondegreen/RoC-ITS). Briefly, the consensus base at any given position of the MSA was determined by the most abundant feature at that position (base or gap). Gaps are then eliminated to produce a consensus sequence.

### Illumina V4 16S libraries

The mock community sample used here (Table 4) was sequenced both with RoC-ITS and with Illumina sequencing-by-synthesis technology. Illumina sequencing was carried out via amplification of the V4 hypervariable region of the 16S rRNA gene with the Earth Microbiome modified F515/R806 primers (22) on a MiSeq platform using a MiSeq 600 cycle kit (Illumina, San Diego, CA, USA). Analysis of the V4 regions was carried out using QIIME2 (21). Amplicon sequencing variants (ASVs) were generated using the DADA2 subcommand (30) from within QIIME2 release 2018.11 with the parameters “--p-trim-left-f 31 --p-trim-left-r 32 -- p-trunc-len-f 220 --p-trunc-len-r 150”. A portion of the Mothur MiSeq SOP (39) was then followed to align reads to the RDP training set v.9 (40) and to remove reads identified as anything other than eubacteria or archaea. Remaining reads were imported back into QIIME2 and chimeras were removed using the “vsearch uchime-denovo- subcommand (41). ASVs were classified using the “classify-sklearn- command in QIIME2 against release 132 off the Silva SSU database (42).

**Table 4.** Mock Community Taxonomy and Metrics.

|  | Phylum | Class | Genus | # of Illumina<br>V4 reads | # of RoC-ITS<br>Consensus<br>Sequences | Mean ITS<br>Length | STD ITS<br>Length | % ID to<br>Consensus | STD to<br>Consensus |
| --- | --- | --- | --- | --- | --- | --- | --- | --- | --- |
| 1 | Bacteroidetes | Bacteroidia | Flavobacterium | 55,508 | 433 | 770.3 | 12.5 | 97.6 | 0.01264 |
| 2 | Actinobacteria | Actinobacteria | - | 38,772 | 135 | 733.4 | 43.4 | 95.3 | 0.01818 |
| 3 | Proteobacteria | Gammaproteobacteria | Pseudomonas | 28,598 | 204 | 615.5 | 124.6 | 97.8 | 0.02182 |
| 4 | Proteobacteria | Gammaproteobacteria | Pseudomonas | 26,951 | 148 | 716.5 | 91.1 | 96.6 | 0.01903 |
| 5 | Firmicutes | Bacilli | Bacillus | 26,521 | 232 | 427.9 | 66.6 | 97.3 | 0.02029 |
| 6 | Proteobacteria | Gammaproteobacteria | Duganella | 12,681 | 80 | 683.5 | 20.2 | 98.1 | 0.01442 |
| 7 | Proteobacteria | Gammaproteobacteria | Pseudomonas | 12,313 | 66 | 708.0 | 146.7 | 97.5 | 0.01702 |
| 8 | Proteobacteria | Gammaproteobacteria | Pseudomonas | 11,697 | 50 | 692.1 | 74.7 | 97.8 | 0.02221 |
| 9 | Bacteroidetes | Bacteroidia | Mucilaginibacter | 9,373 | 87 | 961.7 | 19.7 | 97.5 | 0.017 |
| 10 | Proteobacteria | Alphaproteobacteria | Bradyrhizobium | 4,135 | 33 | 1,160.5 | 29.0 | 97.8 | 0.02157 |
| 11 | Actinobacteria | Actinobacteria | Micrococcus | 3,570 | 9 | 666.5 | 23.2 | 96.5 | 0.03235 |
| 12 | Firmicutes | Bacilli | Bacillus | 3,109 | 95 | 431.1 | 70.9 | 96.5 | 0.02262 |
| 13 | Firmicutes | Bacilli | Bacillus | 2,784 | 36 | 416.8 | 57.7 | 97.1 | 0.02403 |
| 14 | Bacteroidetes | Bacteroidia | Pedobacter | 1,723 | 56 | 1,137.6 | 27.9 | 98.2 | 0.01677 |
| 15 | Proteobacteria | Gammaproteobacteria | Pseudomonas | 1,604 | 4 | 741.8 | 23.7 | 97.9 | 0.0171 |
| 16 | Proteobacteria | Alphaproteobacteria | Inquilinus | 529 | 0 | NA | NA | - | - |
| 17 | Actinobacteria | Actinobacteria | Rhodococcus | 84 | 0 | NA | NA | - | - |
| 18 | Actinobacteria | Actinobacteria | Mycobacterium | 36 | 0 | NA | NA | - | - |

### Analysis of Consensus Sequences

Using cutadapt (version 2.9; (43) the region analogous to the Illumina V4 region was extracted from the RoC-ITS sequences and these were run through QIIME2 to establish a preliminary taxonomic assignment. Not all reads could be assigned by QIIME2 due to indels in the RoC-ITS sequences. To improve the assignment accuracy, the RoC-ITS sequences were aligned to the ASVs from the Illumina V4 regions using NCBI-BLAST (v2.2.26; (44) resulting in the taxonomic assignment of all the RoC-ITS sequences. Chimeric sequences were then identified and removed in an automated fashion with the vsearch software package (version 2.14.2; 34) using a manually curated set of RoC-ITS sequences as the references. The RoC-ITS sequences were then binned by taxonomy, aligned, and corrected with a combination of prank and probcons. The resulting MSAs and phylogenetic trees were manually inspected and a small number of chimeric sequences not detected by the automated approach were removed. The non-chimeric taxonomically binned sequences were then used for all the subsequent phylogenetic analyses. The resulting taxonomic MSAs were also used to generate RoC-ITS consensus sequences. To compute the match identity between any individual RoC-ITS sequence and its corresponding RoC-ITS consensus sequence, mismatches to the ends of the consensus were ignored, and the percent identity was defined as the number of matching bases between two sequences over the number of matched positions, excluding positions that were gaps in both the RoC-ITS consensus and individual RoC-ITS sequence.

### Phylogenetic Tree Generation

Phylogenetic trees were generated from the taxonomically binned RoC-ITS sequences or from 16S-only portion of all the final RoC-ITS sequences. The ITS sequence was removed using cutadapt with the following parameters: -g GGTTACCTTGTTACGACTT --match-read-wildcards -e 0.35 --discard-untrimmed. The 16S-ITS sequences were then multiply aligned with a combination of prank and probcons (see above) to generate an MSA that can be fed to RAxML (v8.2.12; parameters: -f a -# 100 -m GTRCAT -p 12345 -x 12345 -s; 38) to generate a phylogenetic tree with bootstrap values. These trees were visualized on the interactive tree of life web site (iToL; 39). The 16S-only trees included the 16S genes from two archaeal genomes as outgroups: two loci from *Methanotorris igneus* Kol 5 and a single locus from *Nitrosopumilus maritimus* SCM1 (accessions NC_015562 and NC_010085). The eubacterial and archaeal sequences were separately aligned with ssu-align (v 0.1.1; 40) and then masked to identify homologous positions between the archaeal and eubacterial sequences. The resulting alignments were then converted into a tree with bootstrap values using RAxML as above.

## Code availability

Code, recipes, and example data are available on GitHub at: https://github.com/mondegreen/RoC-ITS.

## Data availability

The raw Nanopore sequencing data in fastq format as well as the V4 Illumina MiSeq paired-end reads have been submitted to the NCBI Sequence Read Archive under project PRJNA669399.

## Acknowledgements

We thank the Lennon lab at Indiana University, Bloomington, for providing the mock community, Dr. Irene Newton for her support and suggestions, and the other members of the Center for Genomics and Bioinformatics for their patience and support. We acknowledge support from the IU Pervasive Technology Institute, which is supported by the Lilly Endowment, Inc and by the National Science Foundation under Grant No. CNS-0521433.

---

**Supplemental Figure 1**. Histogram of Nanopore read lengths

**Supplemental Figure 2**. Histogram of inter-splint distances

**Supplemental Figure 3**. Strategy illustrating how RoC-ITS reads with distinct inserts are identified and resolved

**Supplemental Figure 4**. Phylogenetic tree generated from *Duganella* RoC-ITS derived 16S genes (the ITS has been removed) using RAxML with 1000 bootstraps. As in Figure 3, 16S genes from two complete *Duganella* genomes are included. There are no longer well-defined clades so the individual leaves have been labeled with colored circles to indicate their clade membership seen in Figure 3. Black filled circles indicate 90% or greater bootstrap support while open circles indicate that bootstrap support is 15% or less.

