## Supplementary figures and images for "High-Resolution Phylogenetic and Population Genetic Analysis of Microbial Communities with RoC-ITS"

### Supplemental Fig 1

Supplemental Figure 1

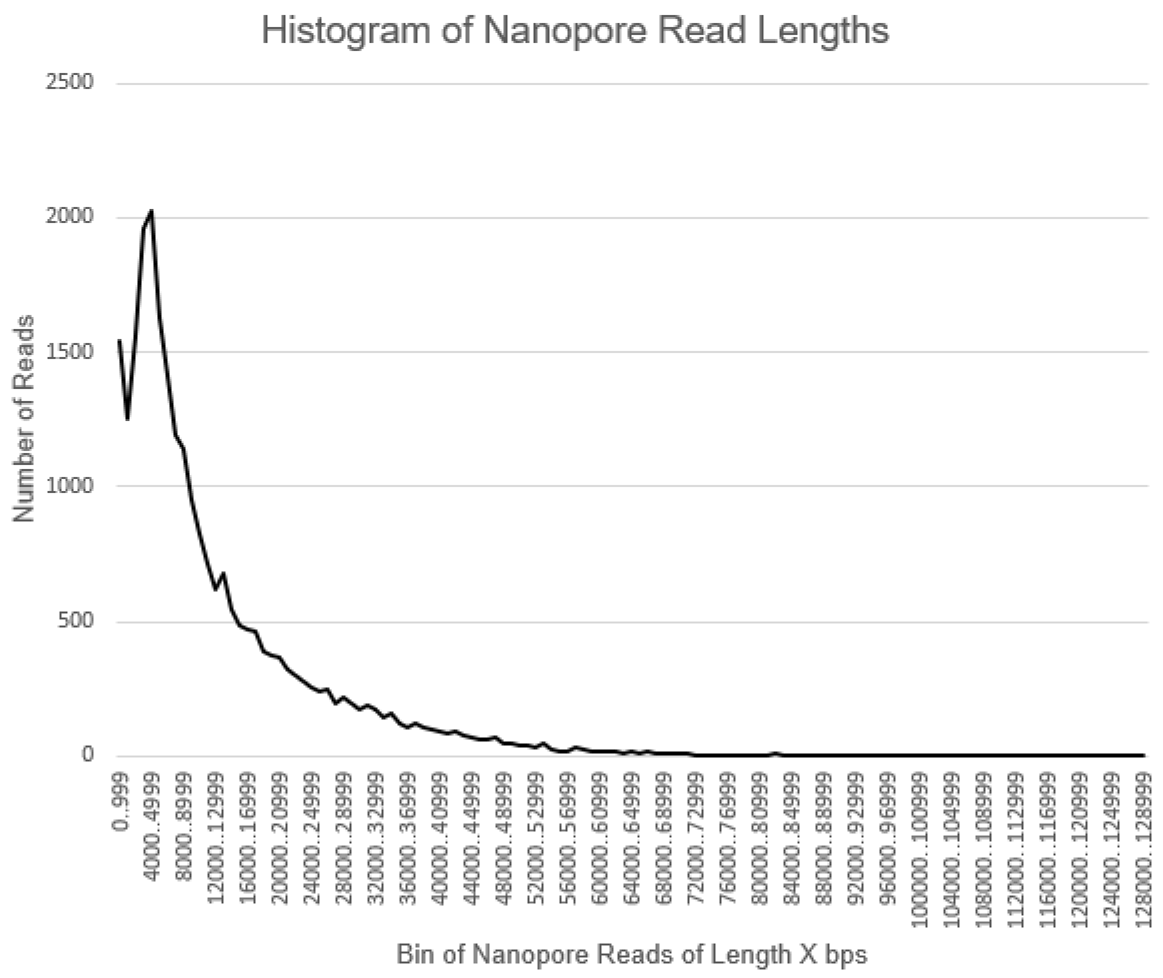

### Supplemental Fig 2

Supplemental Figure 2

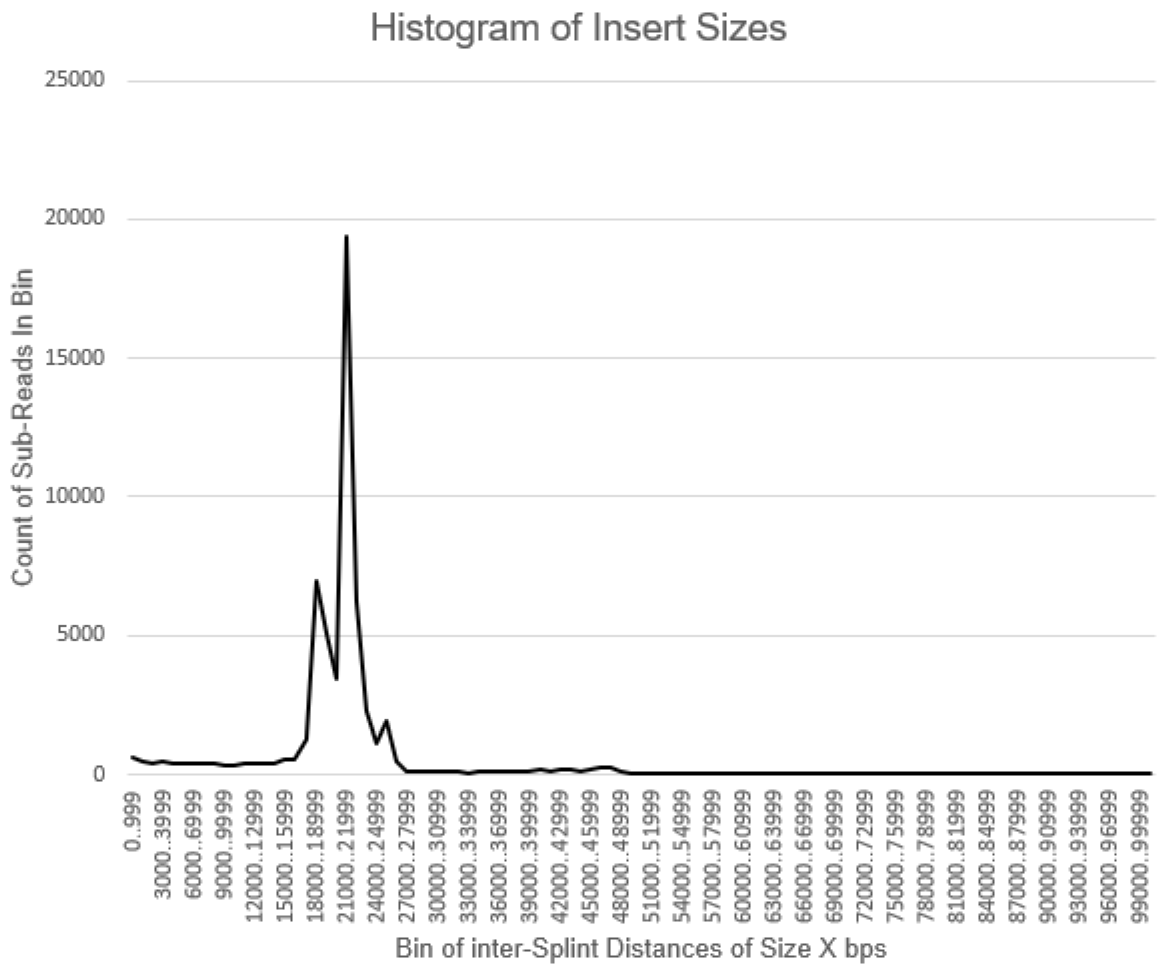

### Supplemental Fig 4

#### Supplemental Figure 4

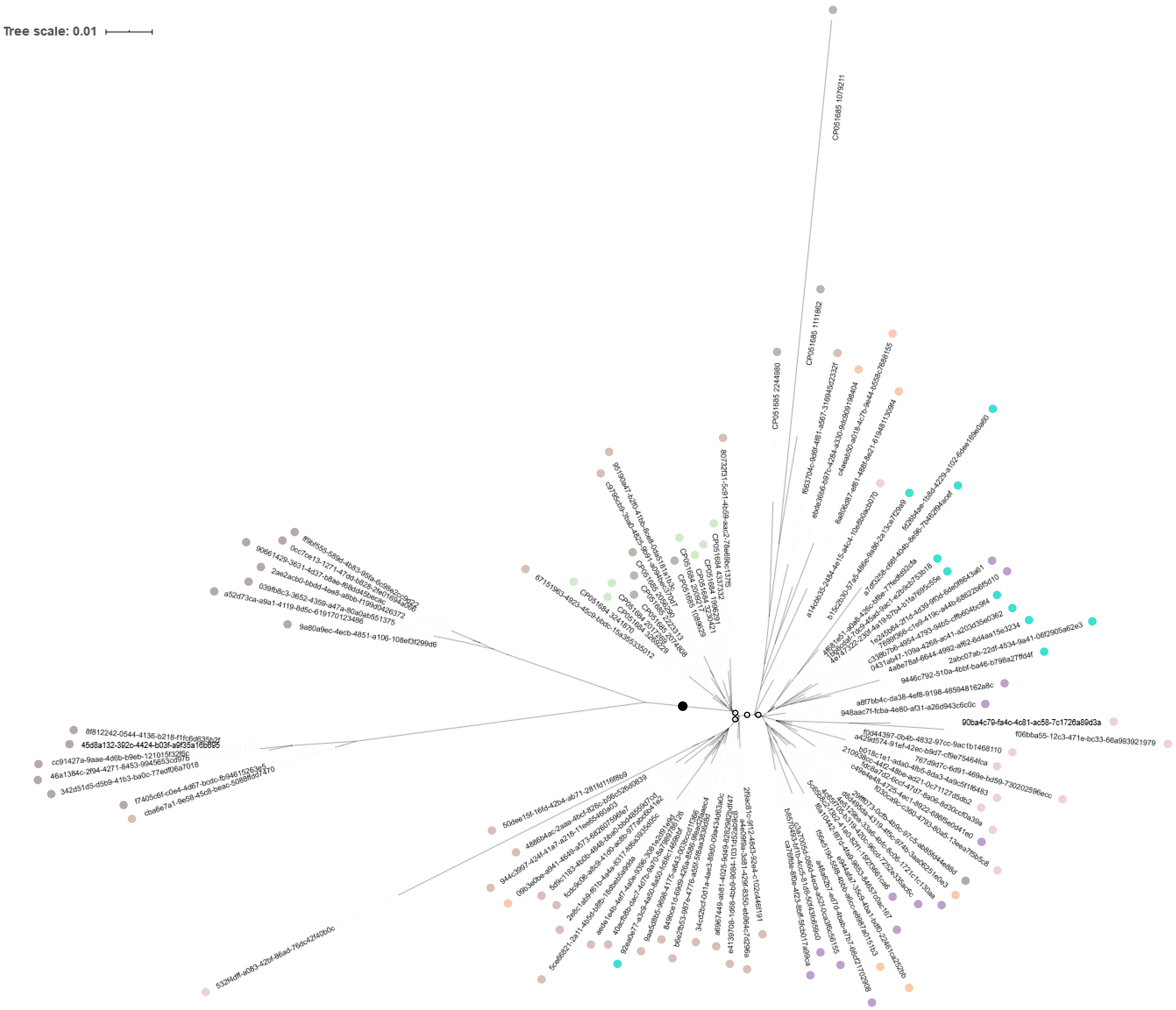
