## Supplemental Fig 3 for "High-Resolution Phylogenetic and Population Genetic Analysis of Microbial Communities with RoC-ITS"

### Supplemental Figure 3

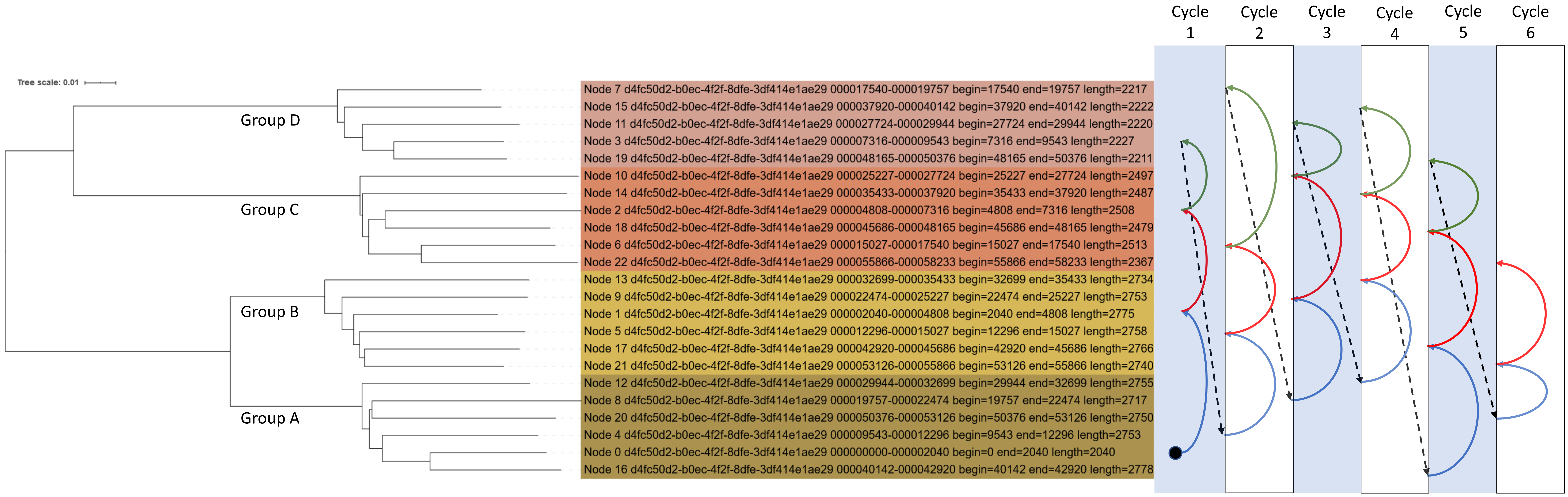


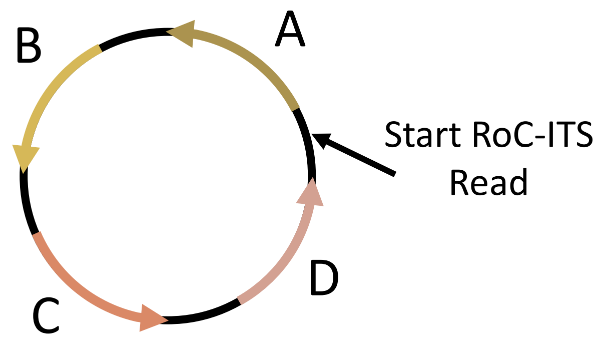


Probabilistic Framework: If there are multiple independent inserts in a single circular template, we expect to see cycles e.g. A→B→C→D→A→B→C→D→A→B→C→D. This can be expressed as a binomial with a 25% probability of success (for example A→A|B|C|D but only B is a success). In this case, with 23 sequences, there are 22 independent trials.
